# Spatial Neuroimmune Crosstalk Driving Perineural Invasion in Head and Neck Squamous Cell Carcinoma

**DOI:** 10.64898/2026.05.26.727931

**Authors:** Riya Chhabra, Alfred Kao, Reena Ding, Suravi Bajaj, Symphony Griffith Jackson, Wei Tse Li, Daniel J. John, Jessica Wang-Rodriguez, Weg M. Ongkeko

## Abstract

**Background:** Perineural invasion (PNI) is a clinically important feature of aggressive head and neck squamous cell carcinoma (HNSCC) and is associated with recurrence and poor survival. However, the spatial organization and molecular programs that characterize tumor–nerve interactions in HNSCC remain incompletely understood.

**Methods:** Single-cell-resolution Xenium spatial transcriptomic data from 10 HNSCC patients were analyzed to define nerve-associated regions, cell-type composition, tumor–nerve proximity, and perineural transcriptional programs. Complementary Visium spatial transcriptomic datasets were used to assess broader transcriptomic features of nerve-associated regions, including epithelial–mesenchymal transition and inferred ligand–receptor signaling. Clinical relevance was evaluated using bulk transcriptomic validation cohorts from TCGA-HNSC and GSE65858.

**Results:** Spatial mapping identified a distinct nerve-proximal microenvironment characterized by depletion of mature dendritic cells and altered immune and stromal composition, consistent with localized disruption of antigen-presenting support at the tumor–nerve interface. HPV− tumors exhibited closer tumor–nerve proximity and a higher exploratory PNI index compared with HPV+ tumors, suggesting subtype-specific differences in nerve-associated tumor behavior. Differential expression analysis of perineural versus distal tumor cells identified a spatially enriched gene program, including NFE2L2, MDM2, and PPARG, that demonstrated a nerve-proximal gradient most evident in HPV− disease. A composite three-gene score was associated with worse overall survival in HPV− patients in TCGA-HNSC and validated in GSE65858, while showing limited prognostic value in HPV+ disease. Visium analyses provided complementary evidence of EMT enrichment and inferred tumor–nerve signaling involving neural guidance and adhesion-associated pathways.

**Conclusions:** These findings support a model in which PNI in HNSCC reflects a spatially organized, transcriptionally distinct tumor–nerve microenvironment, particularly in HPV− disease. The NFE2L2/MDM2/PPARG signature may provide a candidate biomarker for risk stratification and nominates pathways for future mechanistic and therapeutic investigation.

## Introduction

Perineural invasion (PNI)—the infiltration of tumor cells within, around, or through nerve fibers—is a hallmark of aggressive tumor biology and an independent predictor of poor prognosis in head and neck squamous cell carcinoma (HNSCC). Despite its high prevalence, reported in up to 90% of cases [1–4], the molecular and spatial mechanisms enabling PNI remain poorly characterized.

The tumor-nerve interface is not a uniform zone, and detecting transcripts using standard bulk RNA-seq does not fully capture the behavior of individual cell populations [5,6]. Spatial transcriptomics is a relatively recent technology that preserves the tissue context lost in these bulk sequencing approaches. It retains each cell’s position within the tissue, allowing the expression data to be interpreted alongside neighborhood composition and distance relationships. The ability to see the proximity of certain cell types within a tumor can, therefore, provide deeper insight into cellular communication and yield mechanistically plausible pathway explanations [7,8]. This spatial analysis is especially beneficial for investigating PNI, given the inherently distance-dependent nature of tumor-nerve crosstalk and cancer metastasis [9].

Perineural invasion is increasingly recognized as an active interaction between tumor cells and the surrounding neural microenvironment rather than a passive pattern of local spread [10,11]. The perineural niche contains not only tumor cells and nerves, but also Schwann cells, immune populations, endothelial cells, fibroblasts, and extracellular matrix components [2,10,12]. Prior studies have implicated neurotrophic factors such as nerve growth factor and glial cell line-derived neurotrophic factor in tumor cell survival and nerve-directed migration [13]. However, PNI likely reflects a broader spatial program involving immune regulation, stromal remodeling, metabolic adaptation, and cell–cell communication [8,12,14].

Traditional bulk transcriptomic approaches are limited in their ability to resolve these relationships because they average gene expression across mixed cell populations and lose spatial context. Spatial transcriptomics offers an opportunity to examine how tumor cells, immune cells, stromal populations, and nerve-associated cells are organized relative to one another within intact tissue [6,7]. This is particularly important for PNI, where biological behavior is inherently distance-dependent.

The divergent oncogenic biology of HPV-positive and HPV-negative HNSCC provides a critical framework for understanding differential susceptibility to perineural invasion. HPV-negative tumors are predominantly driven by tobacco- and alcohol-related carcinogenesis, frequently harboring TP53 mutations, chromosomal instability, and broad epigenomic dysregulation [15,16]. Loss of p53 function in these tumors not only impairs apoptotic checkpoints but also licenses transcriptional programs associated with stromal remodeling and invasive migration [17,18]. In contrast, HPV-positive tumors are driven by viral oncoproteins E6 and E7, which degrade p53 and pRb through distinct mechanisms [19,20], but generally exhibit less genomic instability and are associated with superior prognosis and treatment response. These mechanistic differences suggest that the capacity for nerve-directed invasion may be more pronounced in HPV-negative disease, rather than representing a uniform feature of HNSCC biology. However, the spatial organization of these interactions and the extent to which they define a distinct invasive tumor state remain poorly understood.

In this study, spatial transcriptomic datasets were integrated to characterize the cellular and molecular architecture of the perineural niche in HNSCC. Using single-cell-resolution Xenium data, nerve-associated regions were defined, tumor–nerve proximity was quantified, and perineural tumor cell programs were identified. Complementary Visium datasets were used to evaluate broader transcriptomic features of nerve-associated regions, including EMT enrichment and inferred ligand–receptor signaling. Finally, spatially derived gene programs were evaluated in independent bulk transcriptomic cohorts to assess their clinical relevance. Together, this approach provides a spatially resolved framework for understanding PNI as an organized tumor–microenvironment state in HNSCC.

## Materials and Methods

### Datasets

Spatial transcriptomic datasets were obtained from publicly available Gene Expression Omnibus (GEO) repositories. Xenium single-cell spatial transcriptomic data were derived from GSE300147. Visium spatial transcriptomic datasets included GSE281978 (six samples; three primary HNSCC tumors and three matched lymph node metastases, of which only primary tumors were analyzed) and GSE181300 (eight samples from two patients, of which four “invasive front” samples were analyzed).

Bulk transcriptomic validation was performed using The Cancer Genome Atlas Head and Neck Squamous Cell Carcinoma cohort (TCGA-HNSC; n = 487 with HPV annotation) and an independent external dataset from GEO (GSE65858; n = 270).

### Xenium Analyses

#### Spatial Transcriptomic Dataset

Xenium spatial transcriptomic data (10x Genomics) from GSE300147 included 10 HNSCC patient samples (5 HPV+ and 5 HPV−), with one ameloblastoma sample excluded. Analyses were performed on Run 1 data following quality control, resulting in a final panel of 399 genes.

#### Quality Control and Preprocessing

Cells with fewer than 5 detected transcripts or fewer than 3 expressed genes were excluded. Gene expression values were normalized to 10,000 counts per cell and log1p-transformed. Batch effects across patients were corrected using Harmony. Dimensionality reduction was performed using principal component analysis (20 components), and clustering was conducted using the Leiden algorithm (resolution = 0.5, k = 15).

#### Spatial PNI Zone Definition

Nerve-associated cells were identified using a composite expression score based on established Schwann cell markers (PMP22, EDNRB, PTN, LGI4) [21-24]. Individual marker scores were calculated using sc.tl.score_genes in Scanpy, and the composite score was defined as the mean of the four marker scores. A threshold of 0.8 was selected empirically based on the composite score distribution and threshold sensitivity analysis. This cutoff captured the high-score tail of cells with elevated nerve-associated marker expression while excluding the majority of low-scoring background cells (Supplementary Figure S1). Sensitivity analysis across candidate cutoffs from 0.4 to 1.2 showed the expected decrease in cell yield with increasing marker-stringency, supporting 0.8 as a conservative cutoff that retained sufficient cells for downstream spatial zone analysis (Supplementary Figure S2).

Spatial zones were defined based on Euclidean distance to the nearest nerve-associated cell using a k-d tree approach, consistent with spatial transcriptomic analyses that use cell coordinates, physical proximity, and local neighborhood structure to characterize tissue microenvironments [25–27]. Cells were categorized into four zones: nerve-associated, perineural (≤7 μm), peritumoral/local microenvironmental (≤20 μm), and distal (>20 μm). The distance thresholds were selected empirically to distinguish cells in immediate proximity to nerve-associated cells from cells in the broader local microenvironment, with ≤7 μm approximating immediate cell-cell proximity and ≤20 μm representing a conservative local neighborhood radius.

#### Cell Type Annotation and Validation

Cells were annotated into nine populations (Tumor, Proliferating Tumor, T cell, B cell, Macrophage, DC_LAMP3, Endothelial, Fibroblast, Mast) using canonical marker gene sets.

Annotations were validated using three complementary approaches:

1. Pearson correlation of mean expression profiles with a published HNSCC single-cell reference dataset from Puram et al. (GSE103322) [28] (Supplementary Figure S3),
2. scoring of the DC_LAMP3 cluster using a curated mature/migratory dendritic cell marker set consisting of LAMP3, CCR7, CD83, CXCL9, CXCL10, and CD274, selected based on published markers of LAMP3^+^/CCR7^+^ activated dendritic cell states and interferon-associated chemokine expression [29,30] (Supplementary Figure S4),
3. marker gene specificity analysis demonstrating enrichment of canonical markers within each annotated population.

#### Neighborhood Analysis

Spatial neighborhood enrichment was assessed using permutation-based analysis. Cells within 50 μm of each nerve-associated cell were identified using scipy cKDTree, with 200 permutations performed to calculate enrichment z-scores. Cell-to-nerve distance distributions were compared between HPV groups using Mann–Whitney U tests.

#### Perineural Invasion Index

A perineural invasion (PNI) index was defined as the ratio of perineural tumor cells to nerve-associated cells for each patient. This metric was designed to quantify tumor infiltration within the immediate neural niche independent of overall nerve-associated cell density and is presented as a pilot measure (n = 10). Given the limited number of patients in the Xenium cohort, patient-level comparisons were interpreted as exploratory and hypothesis-generating. Cell-level analyses were used to characterize spatial organization within the dataset, while independent bulk transcriptomic cohorts were used to assess the clinical relevance of spatially derived gene programs.

For the PNI index comparison (n = 5 per HPV group), per-patient PNI index values were extracted from the Xenium AnnData object by computing the ratio of perineural tumor cells (≤7 μm from nerve-associated cells) to nerve-associated cells per patient. HPV− patients exhibited a median PNI index of 3.33 versus 0.69 in HPV+ patients (Mann-Whitney U test, p = 0.016). The observed rank-biserial correlation |r| = 0.92 indicated a large effect size, consistent with near-complete separation of HPV− and HPV+ PNI index values. Post-hoc power analysis estimated 76% power at α = 0.05 (two-tailed, based on Cohen’s d = 1.92), further supporting the observed large effect size while acknowledging the limited patient number. All spatial findings were treated as hypothesis-generating and were independently validated in bulk transcriptomic cohorts comprising 757 patients across two datasets (TCGA-HNSC, n = 487; GSE65858, n = 270).

#### Differential Gene Expression Analysis

Differential expression was performed to identify genes enriched in tumor cells located near nerve-associated cells. Tumor cells were subset from the Xenium dataset and classified by spatial zone based on distance to the nearest nerve-associated cell. Single-cell differential expression compared perineural tumor cells, defined as tumor cells within ≤7 μm of a nerve-associated cell, against distal tumor cells, defined as tumor cells >20 μm from the nearest nerve-associated cell. Genes were tested using Wilcoxon rank-sum tests, with spatial zone as the grouping variable, followed by false discovery rate correction. Genes with FDR < 0.05 and |log2FC| > 1 were considered significant.

To account for patient-level variability, pseudobulk differential expression was also performed. Raw counts were aggregated by patient and spatial zone among tumor cells, and genes were analyzed using DESeq2 with a design formula of ∼ patient_id + zone. This model tested the effect of spatial zone while accounting for inter-patient variability. Genes with |log2FC| > 0.5 were considered significant. HPV-stratified analyses were then performed separately within HPV− and HPV+ tumors to determine whether nerve-proximal transcriptional programs differed by HPV status.

### Bulk Transcriptomic Validation

#### TCGA Validation

TCGA-HNSC data from the PanCan Atlas 2018 cohort were obtained via cBioPortal. Bulk transcriptomic validation was performed to determine whether the spatially derived perineural tumor-cell gene signature identified in Xenium, composed of NFE2L2, MDM2, and PPARG, was associated with overall survival in an independent patient cohort. For each patient, expression values for NFE2L2, MDM2, and PPARG were individually z-normalized across the cohort. A composite signature score was then calculated as the mean of the three z-normalized expression values:

Composite score = mean(zNFE2L2, zMDM2, zPPARG).

Patients were stratified into signature-high and signature-low groups using the median composite score as the cutoff, which generated balanced groups and avoided arbitrary threshold selection. Survival differences between signature-high and signature-low patients were tested using Kaplan–Meier analysis with a 7-year overall survival endpoint and compared using the log-rank test. Analyses were performed in the full TCGA-HNSC cohort and after stratification by HPV status to determine whether the prognostic association was most evident in HPV− disease. Multivariate Cox proportional hazards models were used to assess whether the composite signature score was independently associated with overall survival after adjustment for age and tumor stage.

#### GSE65858 Independent Validation

Independent validation was performed using GSE65858 (n = 270 HNSCC patients; Illumina HT12v4 microarray). HPV status was derived from hpv16_dna_rna annotations. The three-gene signature was applied using the same methodology as in TCGA, and overall survival was evaluated using Kaplan–Meier analysis with a 5-year endpoint.

### Visium Analyses

#### Preprocessing & Quality Control

Visium datasets were processed in R using Seurat (v5.4.0). Raw data were loaded using Load10X_Spatial, and datasets were merged into a single object. Low-quality spots were filtered using the following criteria: nCount (250–50,000), nFeature (200–7,500), percent mitochondrial reads <15%, and percent ribosomal reads <40%. Normalization was performed using SCTransform.

#### Integration and Clustering

Dimensionality reduction was performed using principal component analysis, followed by unsupervised clustering using the FindNeighbors and FindClusters functions. Batch effects were corrected using Harmony, and clusters were visualized using UMAP.

#### Cell type annotations

Cell types were annotated based on cluster-specific marker genes identified using FindMarkers. Nerve-associated spots were identified using UCell scoring of canonical neuronal and Schwann cell markers (S100B, SOX10, MPZ). Spots with a UCell score > 0.1 were classified as nerve-associated.

#### Gene Set Signature Scoring

Gene set enrichment for epithelial–mesenchymal transition (EMT) and perineural invasion (PNI) signatures was calculated using the UCell package, which applies a rank-based scoring method based on the Mann–Whitney U statistic. Canonical gene sets were curated from published literature. The EMT signature was constructed using established EMT-associated markers, including VIM, CDH2, FN1, SNAI1, and TWIST1 [31]. PNI-associated gene sets, including nerve markers, neurotrophic factors, axon guidance factors, matrix metalloproteinases, and adhesion factors, are listed in Supplementary Table S1.

#### Ligand-Receptor cell communication inference

Cell–cell communication was inferred using SpatialCellChat (CellChat v3), incorporating spatial constraints. Three tissue samples with a robust subset of nerve-associated cells (n>50) were selected for cell-cell communication analysis. Interaction distances were defined based on spot resolution (∼55 μm diameter), with cell–cell contact interactions restricted to ≤100 μm and secreted or extracellular matrix signaling evaluated within ≤200 μm following the SpatialCellChat vignette. Significant ligand–receptor interactions were identified using permutation-based testing and visualized using circle and bubble plots.

### Data Availability

The data analyzed in this study were obtained from publicly available repositories. Xenium spatial transcriptomic data were obtained from Gene Expression Omnibus (GEO) at GSE300147. Visium spatial transcriptomic datasets were obtained from GEO at GSE281978 and GSE181300. Bulk transcriptomic validation data were obtained from TCGA-HNSC via cBioPortal and from GEO at GSE65858. Derived data supporting the findings of this study are available from the corresponding author upon reasonable request.

### Code Availability

Custom code used for spatial preprocessing, cell-type annotation validation, nerve-associated cell scoring, spatial zone assignment, differential expression analysis, signature scoring, survival analysis, and figure generation is available from the corresponding author upon reasonable request.

## Results

### Spatial transcriptomics reveals organized neuroimmune architecture in HNSCC

To define the cellular composition and spatial organization of the tumor microenvironment, we analyzed single-cell spatial transcriptomic data comprising 560,492 cells across 10 HNSCC patient samples. Unsupervised clustering identified nine major cell populations, including tumor, proliferating tumor, macrophages, T cells, B cells, endothelial cells, fibroblasts, dendritic cells, and mast cells (Fig. 1A–E).

**Figure 1.**
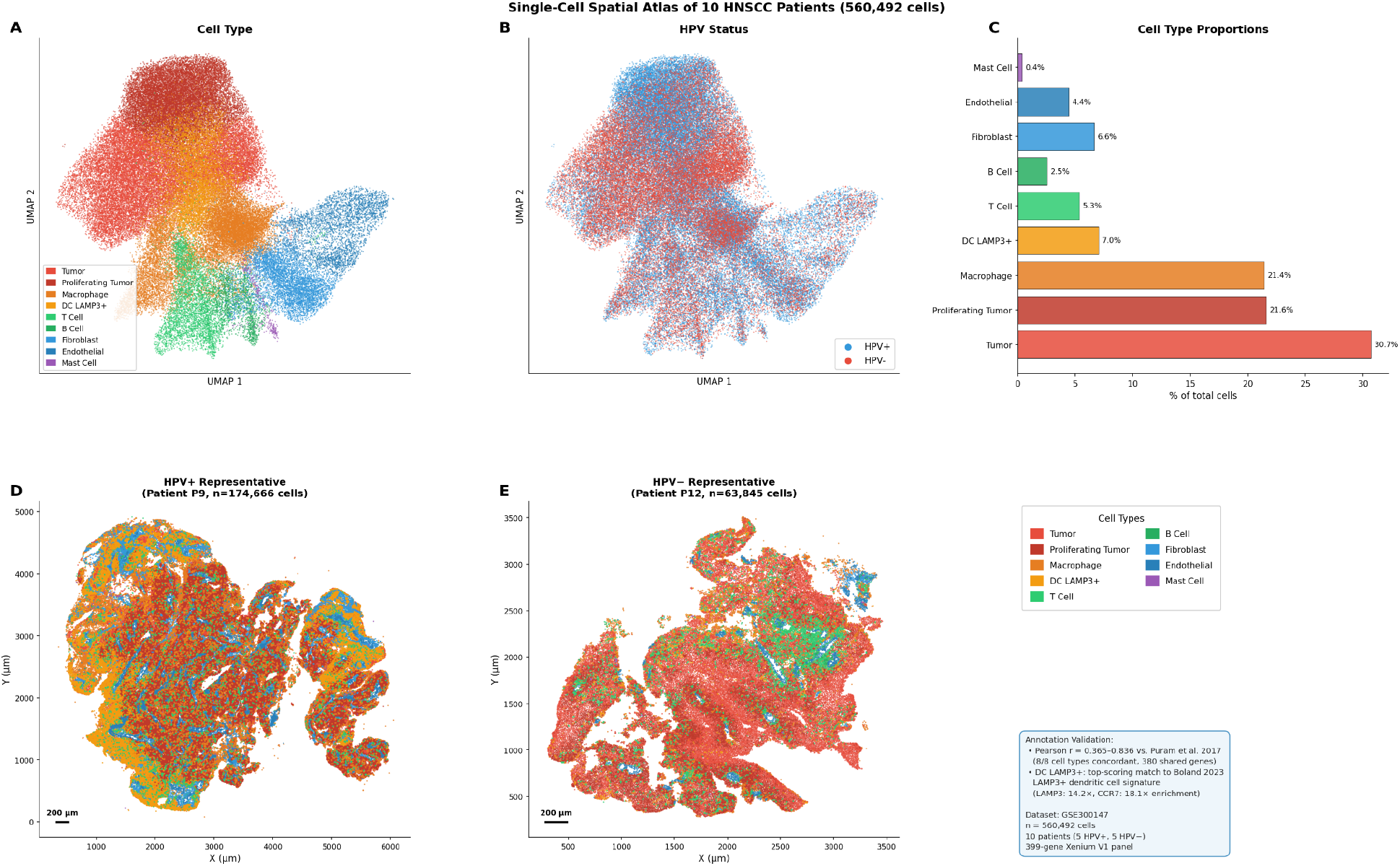
Single-cell spatial atlas of HNSCC tumors profiled by Xenium. (A) UMAP visualization of 560,492 cells from 10 HNSCC patient samples colored by annotated cell type, including tumor, proliferating tumor, macrophage, DC_LAMP3, T cell, B cell, endothelial, fibroblast, and mast cell populations. (B) UMAP visualization colored by HPV status, demonstrating representation of both HPV+ and HPV− tumors within the integrated dataset. (C) Bar plot showing the relative abundance of major annotated cell populations across the full Xenium cohort. Tumor and proliferating tumor cells comprised the largest fraction of cells, followed by macrophages and other immune and stromal populations. (D–E) Representative spatial maps from HPV+ and HPV− tumors showing the spatial distribution of annotated cell populations within intact tissue architecture.

Tumor (29%) and proliferating tumor cells (22%) comprised over half of all cells, alongside substantial macrophage infiltration (21%), consistent with an immunologically active tumor microenvironment. Cell type annotations demonstrated concordance with established reference datasets and canonical marker profiles, supporting classification of tumor, immune, and stromal populations (Supplementary Figures S3–S4).

Together, these data establish the cellular composition and spatial organization of the Xenium HNSCC cohort and support downstream analysis of tumor, immune, stromal, and nerve-associated spatial relationships.

### Spatial framework defines a perineural niche with selective immune dysfunction

To investigate tumor–nerve interactions, nerve-associated cells were identified based on canonical Schwann cell markers (PMP22, EDNRB, PTN, LGI4), and cells were stratified into four spatial zones: nerve-associated, perineural (≤7 μm), peritumoral (≤20 μm), and distal (>20 μm). This spatial framework enabled systematic analysis of cellular composition as a function of distance from nerve-associated cells (Fig. 2A–C).

**Figure 2.**
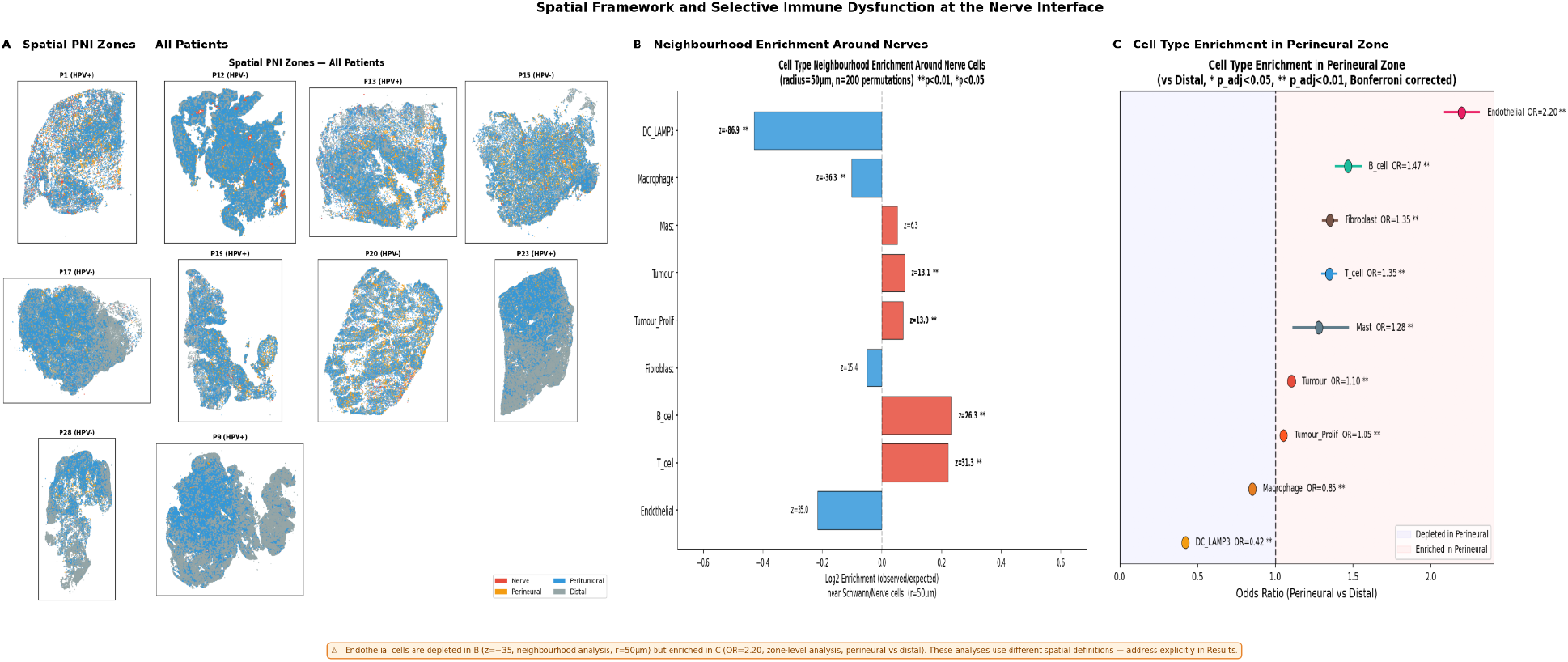
Spatial framework defines nerve-associated regions and reveals selective immune remodeling near Schwann/neural marker-expressing cells. (A) Representative spatial maps showing the classification of cells into spatial zones based on Euclidean distance to the nearest Schwann/neural marker-expressing nerve-associated cell. Zones included nerve-associated cells, perineural cells located within ≤7 μm, peritumoral/local microenvironmental cells located within ≤20 μm, and distal cells located >20 μm from the nearest nerve-associated cell. (B) Neighborhood enrichment analysis showing cell-type enrichment or depletion within 50 μm of nerve-associated cells across 10 Xenium HNSCC samples. DC_LAMP3+ mature dendritic cells showed the strongest depletion near nerve-associated cells, while lymphocyte populations showed relative enrichment. (C) Odds ratio analysis comparing cell-type representation in the perineural zone versus distal regions across 10 Xenium HNSCC samples. Mature dendritic cells were underrepresented in the perineural zone, while endothelial cells were enriched, supporting spatially selective remodeling of the nerve-proximal microenvironment.

Using this framework, spatial analysis revealed a highly organized immune architecture surrounding Schwann/neural marker-expressing nerve-associated cells. DC_LAMP3^+^ served as a marker for mature dendritic cells (mDCs), which were the most significantly depleted population near nerve-associated cells (z = −88), followed by macrophages (z = −39), indicating a marked reduction in antigen-presenting capacity within the perineural niche. In contrast, both T cells and B cells were enriched near nerve-associated cells, indicating that immune cells are present but spatially uncoupled from antigen-presenting support.

Odds ratio analysis confirmed these findings, with mDCs significantly underrepresented in the perineural zone (OR = 0.42), while endothelial cells were enriched (OR = 2.20). Notably, the apparent discrepancy between endothelial depletion in neighborhood enrichment and enrichment in odds ratio analysis likely reflects differences in spatial scale, where local proximity-based analyses capture immediate cellular neighborhoods, while compositional analyses reflect broader regional abundance.

Together, these findings suggest that the nerve-proximal niche is not uniformly immune-depleted, but instead selectively altered in a way that may reduce local antigen-presenting support. The marked depletion of DC_LAMP3 mature dendritic cells, together with enrichment of T cells and B cells, indicates that lymphocytes are present near nerve-associated cells but spatially uncoupled from a major antigen-presenting population. This pattern defines a localized immune-dysfunctional niche at the tumor–nerve-associated cell interface, characterized by preserved lymphocyte presence but reduced antigen-presenting support.

Together, these analyses identify a localized immune-dysfunctional niche surrounding Schwann/neural marker-expressing nerve-associated cells, characterized by reduced antigen-presenting cell support despite the presence of lymphocytes.

### HPV− tumors exhibit enhanced nerve proximity and increased perineural invasion

We next assessed whether tumor–nerve interactions differed by HPV status. Across multiple cell types, HPV− tumors exhibited significantly closer proximity to nerve-associated cells compared to HPV+ tumors. Tumor cells in HPV− samples were a median of 4.8 μm closer to nerve-associated cells (11.0 μm vs 15.8 μm; Mann–Whitney U test, p < 0.0001), indicating a consistent shift toward nerve-associated localization (Fig. 3A).

**Figure 3.**
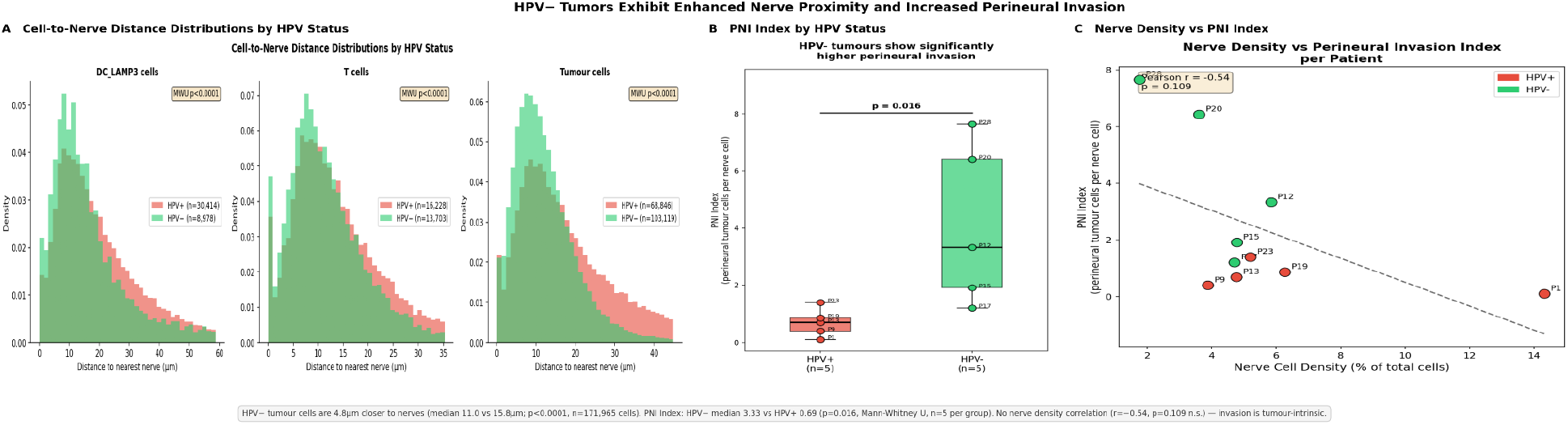
HPV− tumors exhibit increased tumor proximity to nerve-associated cells and a higher exploratory perineural invasion index. (A) Distributions of cell-to-nerve-associated-cell distances stratified by HPV status across major cell populations in 5 HPV+ and 5 HPV− Xenium HNSCC samples. Tumor cells in HPV− samples were located significantly closer to nerve-associated cells than tumor cells in HPV+ samples, suggesting HPV-associated differences in spatial tumor–nerve organization. (B) Per-patient perineural invasion index, defined as the ratio of perineural tumor cells within ≤7 μm of nerve-associated cells to the number of nerve-associated cells, across 5 HPV+ and 5 HPV− samples. HPV− tumors demonstrated a higher PNI index compared with HPV+ tumors, consistent with increased tumor enrichment near nerve-associated regions. (C) Correlation analysis comparing nerve-associated cell density with PNI index across 10 Xenium HNSCC samples. The lack of a significant positive correlation suggests that increased PNI index in HPV− tumors is not simply explained by greater nerve-associated cell abundance.

To quantify perineural invasion, we defined a PNI index as the ratio of perineural tumor cells to nerve-associated cells. HPV− tumors exhibited a greater than four-fold higher PNI index compared to HPV+ tumors (median 3.33 vs 0.69; p = 0.016; Fig. 3B). This difference was not explained by nerve-associated cell density (r = −0.54, p = 0.109; Fig. 3C), suggesting that enhanced tumor localization near nerve-associated regions was not simply attributable to greater nerve-associated cell abundance. Notably, one HPV+ patient (P9) exhibited a PNI index of 2.50, overlapping with the lower range of HPV− values, consistent with the biological heterogeneity observed within HPV+ disease; however, this patient had the highest nerve-associated cell density of the HPV+ cohort, suggesting that elevated nerve-associated cell abundance rather than tumor-intrinsic invasive behavior may contribute to this observation.

These findings identify HPV status as an important determinant of tumor–nerve-associated cell organization and suggest that HPV− tumors preferentially engage nerve-proximal invasive behavior.

These results support HPV status as a determinant of tumor localization near nerve-associated cells and suggest that HPV− tumors demonstrate enhanced nerve-proximal tumor behavior independent of nerve-associated cell density.

### Tumor cells proximal to nerve-associated cells activate a perineural gene program

Differential expression analysis comparing tumor cells in perineural versus distal regions identified a conserved transcriptional program associated with nerve proximity. A total of 43 genes were upregulated and 25 downregulated, with 23 genes concordant across single-cell and pseudobulk analyses, defining a high-confidence perineural gene set.

Among the concordant genes, NFE2L2, MDM2, and PPARG represented a biologically coherent axis involving oxidative stress adaptation, p53 pathway regulation, and metabolic reprogramming [32–35]. This program exhibited a clear spatial gradient in HPV− tumors, with highest expression near nerve-associated cells and progressive decline with increasing distance. In contrast, no significant spatial gradient was detected in HPV+ tumors (Kruskal-Wallis test across spatial zones, p = 1.0), indicating uniform expression regardless of nerve proximity and consistent with the absence of an active perineural transcriptional program in HPV+ disease.

These findings identify a conserved, spatially restricted transcriptional program that is most evident in HPV− disease and associated with perineural invasion (Fig. 4A–C).

**Figure 4.**
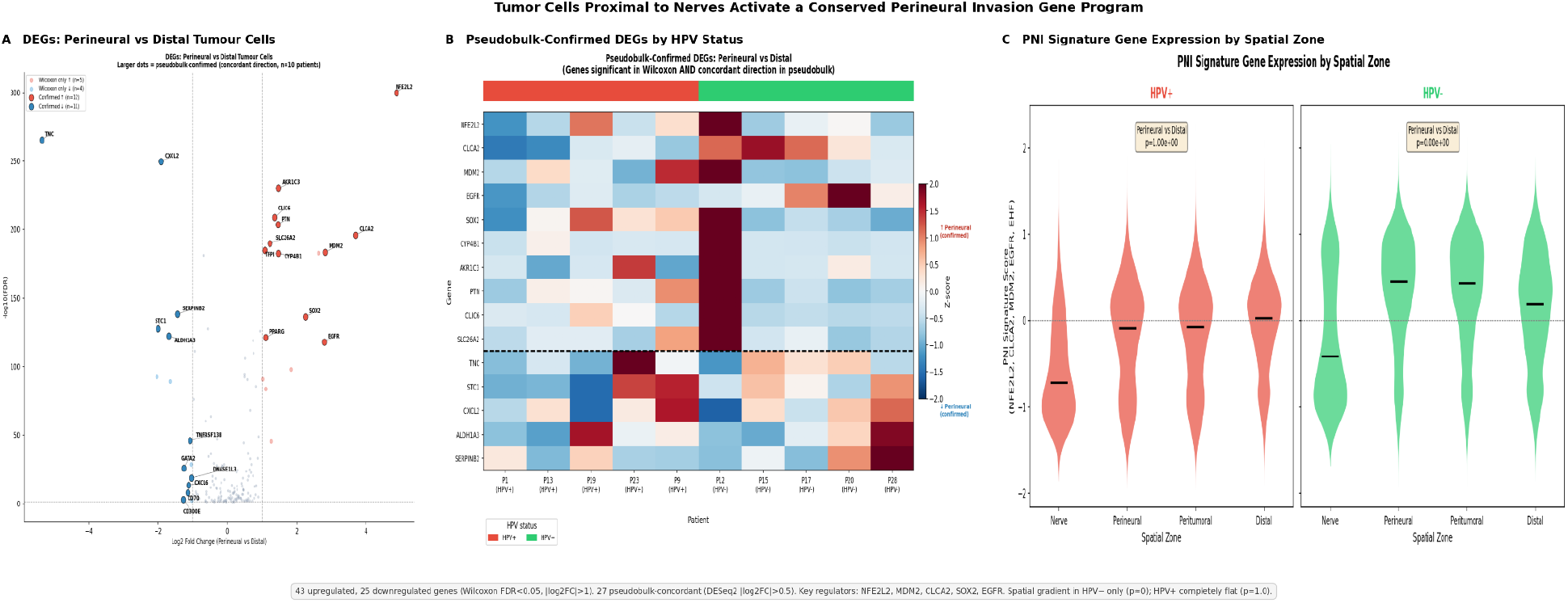
Tumor cells proximal to nerve-associated cells display a spatially enriched perineural transcriptional program. (A) Volcano plot showing differentially expressed genes between perineural tumor cells, defined as tumor cells within ≤7 μm of nerve-associated cells, and distal tumor cells located >20 μm from nerve-associated cells across 10 Xenium HNSCC samples. Differential expression was assessed using single-cell analysis with FDR correction. (B) Heatmap of genes concordant across single-cell and patient-level pseudobulk differential expression analyses, stratified by HPV status. Concordant genes define a high-confidence nerve-proximal tumor-cell transcriptional program. (C) Spatial gradient analysis of the NFE2L2/MDM2/PPARG signature across nerve-associated, perineural, peritumoral, and distal zones. HPV− tumors showed enrichment of the signature near nerve-associated regions with progressive decline across increasing distance, whereas HPV+ tumors did not demonstrate a comparable spatial gradient.

Together, these findings identify a spatially restricted tumor-cell program associated with proximity to nerve-associated cells, most evident in HPV− disease.

### Visium spatial profiling supports EMT enrichment and tumor–nerve ligand–receptor crosstalk

To complement the single-cell-resolution Xenium analysis, we analyzed seven HNSCC tissue sections profiled by Visium spatial transcriptomics. Nerve-associated spots were identified using UCell scoring of canonical Schwann/neural markers (S100B, SOX10, MPZ), yielding 387 nerve-associated spots among 10,047 total spots.

PNI-high spots exhibited enrichment of epithelial–mesenchymal transition (EMT) signatures compared to PNI-low spots (p-value < 2.2e-16), indicating that nerve-proximal regions are associated with increased cellular plasticity and invasive potential. This finding provides independent spatial support for the perineural transcriptional program identified in the Xenium dataset.

SpatialCellChat analysis identified bidirectional ligand–receptor signaling between tumor and nerve-associated cells, including pathways involved in neural development and tumor invasion such as WNT, ephrin, semaphorin, and adhesion signaling. These results support a model in which tumor cells co-opt neural guidance programs to facilitate nerve-directed migration.

Together, these findings provide complementary evidence that perineural invasion is associated with EMT activation and active tumor–nerve signaling (Fig. 5A–C).

**Figure 5.**
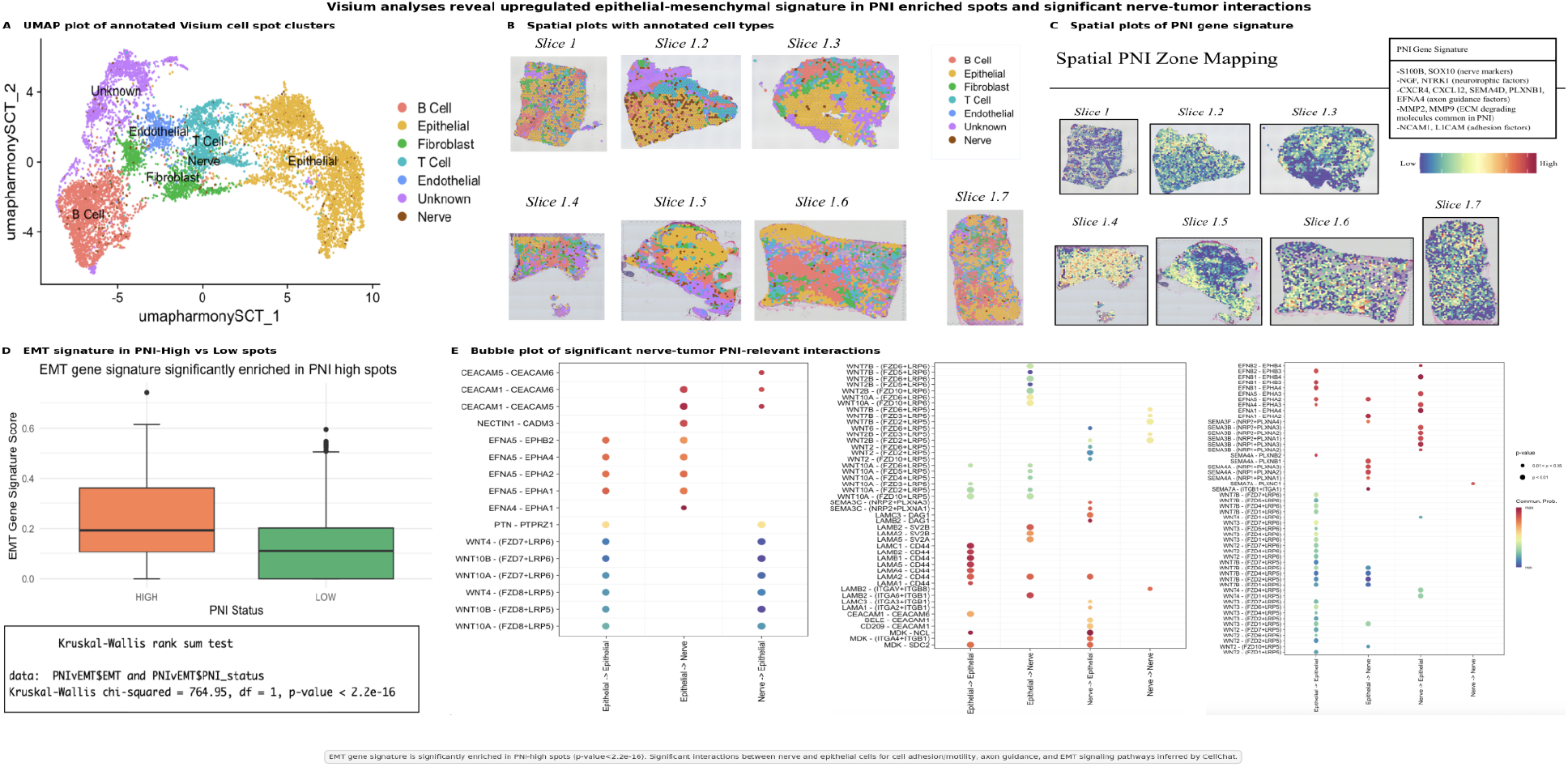
Visium spatial profiling supports EMT enrichment and tumor–nerve-associated signaling. (A) UMAP and representative Visium spatial maps showing annotated cell types and spatial localization of nerve-associated regions across seven HNSCC tissue sections. Nerve-associated spots were identified using UCell scoring of canonical Schwann/neural markers. (B) Comparison of epithelial–mesenchymal transition (EMT) signature scores between PNI-high and PNI-low Visium spots. PNI-high spots showed significantly increased EMT signature enrichment, supporting an association between nerve-proximal regions and invasive cellular plasticity. (C) SpatialCellChat ligand–receptor analysis showing inferred interactions between tumor and nerve-associated cell populations in three tissue sections with a robust subset of nerve-associated cells.

Enriched signaling included neural guidance, adhesion, and tumor invasion-associated pathways, including WNT, ephrin, and semaphorin-related interactions.

These Visium analyses provide complementary transcriptome-wide support for EMT activation and active tumor–nerve-associated signaling in regions enriched for PNI-related features.

### A spatially derived gene signature predicts poor survival in HPV− HNSCC

To evaluate clinical relevance, we applied the perineural gene program to bulk transcriptomic datasets. A three-gene signature (NFE2L2, MDM2, PPARG) stratified HPV− patients into high- and low-risk groups with significantly different overall survival in TCGA (p = 0.0021), while no association was observed in HPV+ patients (p = 0.897).

Multivariate Cox regression confirmed that this signature independently predicted survival after adjustment for clinical variables (HR = 1.29, 95% CI 1.11–1.50, p = 0.0009), and this association was validated in an independent cohort (GSE65858; p = 0.032).

Together, these results demonstrate that a spatially derived nerve-proximal transcriptional program can be translated into a bulk transcriptomic survival signature with strongest prognostic relevance in HPV− HNSCC (Fig. 6A–C).

**Figure 6.**
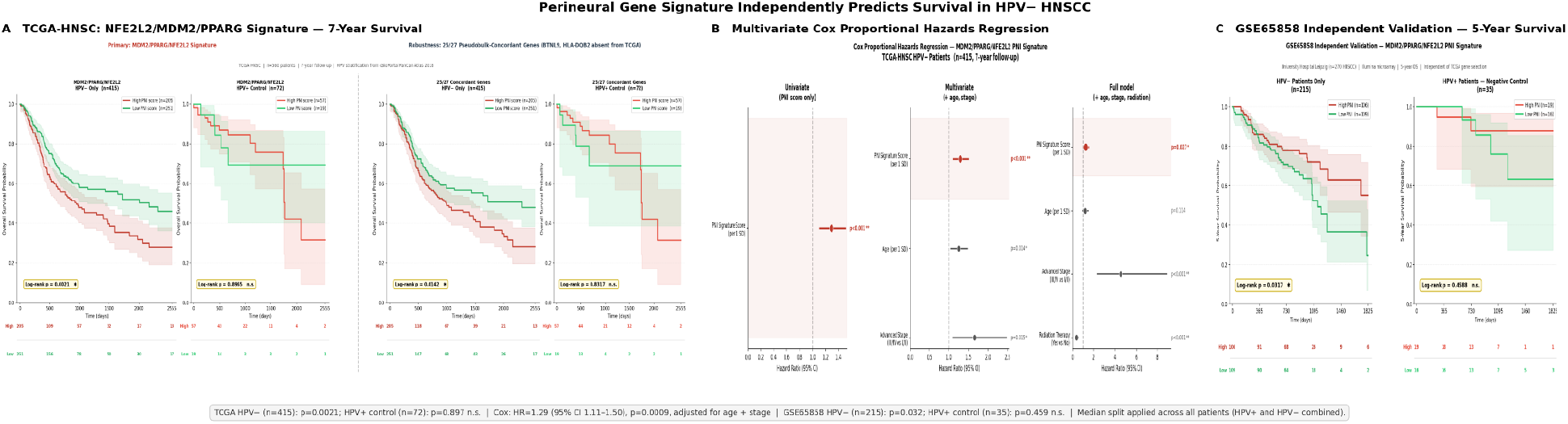
Spatially derived NFE2L2/MDM2/PPARG signature predicts worse survival in HPV− HNSCC. (A) Kaplan–Meier overall survival analysis in TCGA-HNSC using a composite three-gene signature score based on z-normalized expression of NFE2L2, MDM2, and PPARG. Patients were stratified into signature-high and signature-low groups using the median composite score, with analyses performed in the full cohort and after HPV stratification. TCGA-HNSC included 487 patients with HPV annotation. The signature was associated with worse overall survival in HPV− patients but showed limited prognostic value in HPV+ patients. (B) Multivariate Cox proportional hazards analysis assessing whether the composite signature score was independently associated with overall survival after adjustment for clinical covariates, including age and tumor stage. (C) Independent validation in GSE65858 using the same three-gene signature scoring approach and Kaplan–Meier survival analysis with a 5-year endpoint. GSE65858 included 270 HNSCC patients.

Together, these analyses demonstrate that a spatially derived nerve-proximal tumor-cell gene program can be translated into a bulk transcriptomic prognostic signature with strongest relevance in HPV− HNSCC.

## Discussion

This study provides a spatially resolved characterization of the perineural invasion (PNI) niche in head and neck squamous cell carcinoma (HNSCC), integrating single-cell spatial transcriptomics, neighborhood analysis, differential gene expression, ligand–receptor inference, and bulk transcriptomic validation. Across complementary spatial platforms and independent clinical cohorts, these findings support a model in which PNI reflects a spatially organized tumor–microenvironment state, most pronounced in HPV− disease, characterized by altered immune organization, nerve-proximal tumor enrichment, and a clinically relevant transcriptional program.

A central finding of this study is that the perineural niche is not simply immune-depleted, but spatially immune-dysfunctional. Mature dendritic cells were selectively depleted near Schwann/neural marker-expressing nerve-associated cells despite enrichment of T and B lymphocytes, suggesting that immune dysfunction in this niche may arise from spatial uncoupling between lymphocytes and antigen-presenting support rather than global immune exclusion. This organization may create a localized zone of impaired immune surveillance at the tumor–nerve-associated cell interface, allowing tumor cells to persist and invade within an otherwise immune-populated microenvironment.

Several mechanisms may contribute to this spatial pattern. Semaphorin-3A (SEMA3A), which was identified among the ligand-receptor interactions in our SpatialCellChat analysis, has been shown to repel dendritic cell migration through neuropilin-1 signaling, providing a plausible molecular mechanism for DC exclusion at the tumor-nerve interface [36,37]. Additionally, tumor-derived PGE2 and adenosine, both enriched in hypoxic and metabolically active microenvironments, are well-established suppressors of DC maturation and migration [38–40]. The enrichment of PPARG — a known downstream effector of lipid mediator signaling — in perineural tumor cells raises the possibility that metabolic crosstalk between tumor and stromal cells actively shapes the immunological character of the perineural niche [33]. Although these mechanisms remain hypothesis-generating, they suggest that the perineural niche may be actively maintained through coordinated tumor, immune, stromal, and nerve-associated signaling rather than representing a passive consequence of local tumor spread.

HPV status emerged as an important determinant of tumor–nerve-associated cell organization. HPV− tumors demonstrated greater tumor enrichment near nerve-associated regions independent of nerve-associated cell density, supporting the possibility that HPV− tumors are more likely to engage nerve-proximal invasive programs. This distinction is biologically plausible given the divergent oncogenic landscapes of HPV− and HPV+ HNSCC. HPV− tumors more frequently exhibit TP53 disruption, genomic instability, stromal remodeling, and invasive transcriptional states, whereas HPV+ tumors are driven by viral oncogene-mediated pathways and often demonstrate more favorable clinical behavior. The perineural gene program identified in this study, anchored by NFE2L2, MDM2, and PPARG, aligns closely with these differences. NFE2L2, MDM2, and PPARG are linked to oxidative stress adaptation, p53 pathway regulation, and metabolic plasticity, respectively, providing a biologically plausible basis for their enrichment in nerve-proximal HPV− tumor cells [32–35]. Their spatial enrichment near nerve-associated cells suggests that these pathways collectively support tumor cell survival and migration within the perineural microenvironment.

The translation of this spatially derived program into bulk transcriptomic cohorts highlights the clinical relevance of nerve-proximal tumor biology. The NFE2L2/MDM2/PPARG signature was associated with worse survival in HPV− disease and retained prognostic value after adjustment for age and tumor stage, suggesting that spatially defined invasive programs can capture clinically meaningful risk not fully reflected by standard clinicopathologic variables. Because PNI is currently assessed primarily through histopathologic review, a transcriptomic surrogate of nerve-proximal invasive biology could help identify high-risk HPV− tumors when spatial or histologic assessment is limited. These findings support the broader concept that PNI is not only a histopathologic observation, but may also represent a biologically encoded tumor state with measurable transcriptional outputs.

Complementary Visium analyses further reinforced this model by linking nerve-associated regions to epithelial–mesenchymal transition enrichment and inferred ligand–receptor interactions involving WNT, ephrin, semaphorin, and adhesion-associated pathways. Together, these findings suggest that PNI reflects active tumor–microenvironment communication rather than passive local extension. From a therapeutic perspective, the pathways nominated by the spatial gene program, including NRF2 signaling,

MDM2-mediated p53 regulation, and PPARG-associated metabolic plasticity, represent biologically plausible targets for future investigation. NRF2 pathway activity can be modulated by small-molecule inhibitors such as ML385 and brusatol, MDM2 antagonists such as nutlin-3a and AMG-232 have been developed to restore p53 pathway activity in selected contexts, and PPARG inverse agonists such as T0070907 and SR10221 have been explored in models of invasive epithelial cancer [34,35,41–43]. These targets should be interpreted as hypothesis-generating rather than immediately practice-changing, but they provide a rationale for testing whether disruption of oxidative stress tolerance, apoptotic resistance, metabolic adaptation, or neuroimmune signaling can limit tumor–nerve interactions.

Several limitations should be considered. The Xenium platform uses a targeted 399-gene panel, which constrains transcriptome-wide discovery; however, the consistency of findings across independent spatial and bulk datasets supports the robustness of the identified perineural program. Nerve-associated cell identification was based on a composite expression score derived from established Schwann/neural markers rather than direct histologic nerve annotation, which may introduce some degree of misclassification, particularly in regions of low nerve-associated cell density. The Xenium and Visium analyses were also performed on independent cohorts with different tissue processing workflows and gene coverage, so findings across platforms should be interpreted as complementary rather than direct cross-validation. Finally, the spatial discovery cohort was limited to 10 patients, and the PNI index should be regarded as an exploratory metric requiring validation in larger spatial cohorts with matched pathologist-scored PNI status. Functional studies will also be required to establish causal mechanisms underlying the tumor–nerve-associated interactions identified here.

Overall, this study reframes PNI in HNSCC as a spatially organized and transcriptionally measurable tumor–microenvironment state. By identifying immune dysfunction, HPV-associated nerve-proximal tumor behavior, and a prognostic NFE2L2/MDM2/PPARG signature, these findings provide a framework for future studies of PNI biology and nominate candidate biomarkers and pathways for risk stratification and therapeutic investigation in HPV− HNSCC.

## Acknowledgments

Generative artificial intelligence tools were used to assist with language editing, organization, and manuscript formatting. All analyses, scientific interpretations, and final manuscript content were reviewed and approved by the authors. No specific funding was received for this work.

