## Supplementary Figures S1-S4 for "Spatial Neuroimmune Crosstalk Driving Perineural Invasion in Head and Neck Squamous Cell Carcinoma"

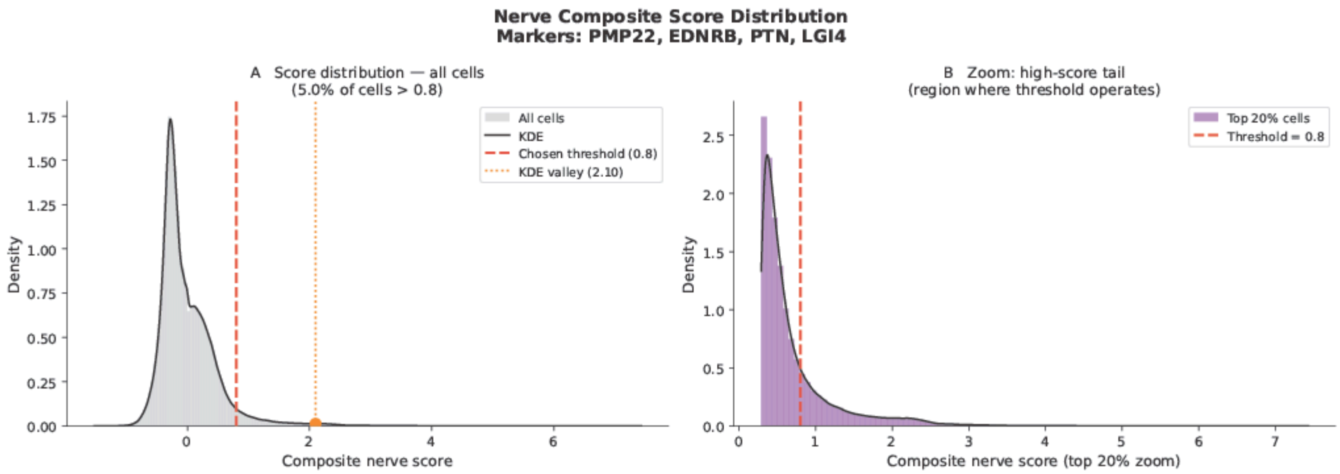


**Supplementary Figure S1. Distribution of nerve-associated composite scores used to identify Schwann/neural marker-expressing cells.**

Composite nerve-associated scores were calculated using expression of PMP22, EDNRB, PTN, and LGI4. The full score distribution across all cells is shown alongside a zoomed view of the high-score tail. A threshold of 0.8 was used to identify cells with elevated Schwann/neural marker expression while excluding the majority of low-scoring background cells.

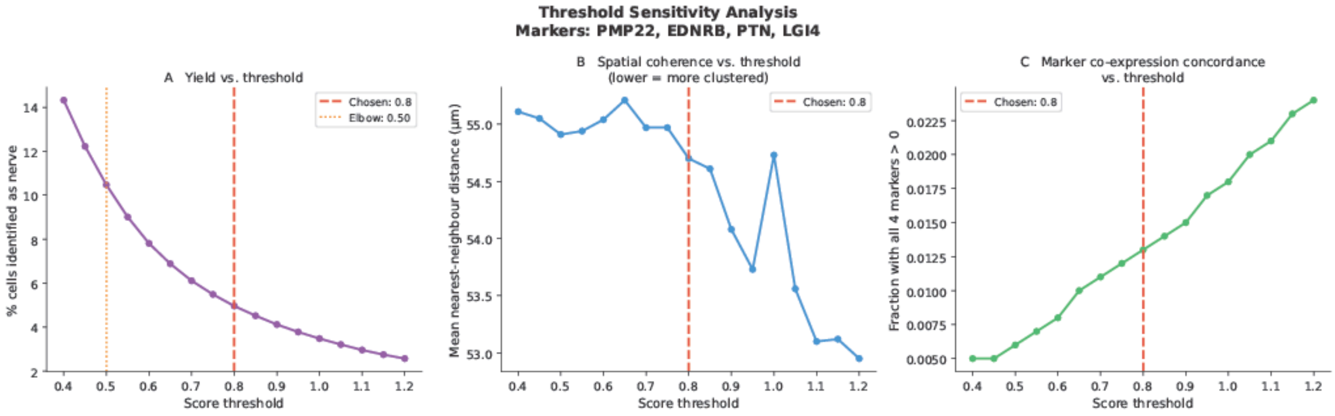


**Supplementary Figure S2. Threshold sensitivity analysis for marker-defined nerve-associated cell identification.**

Sensitivity analysis was performed across candidate composite score thresholds from 0.4 to 1.2. Increasing thresholds reduced the number of cells classified as nerve-associated while increasing marker stringency. When spatial coordinates were available, mean nearest-neighbor distance among selected cells was also calculated as a measure of spatial coherence. The selected threshold of 0.8 balanced conservative marker enrichment with sufficient cell yield for downstream spatial zone and neighborhood analyses.

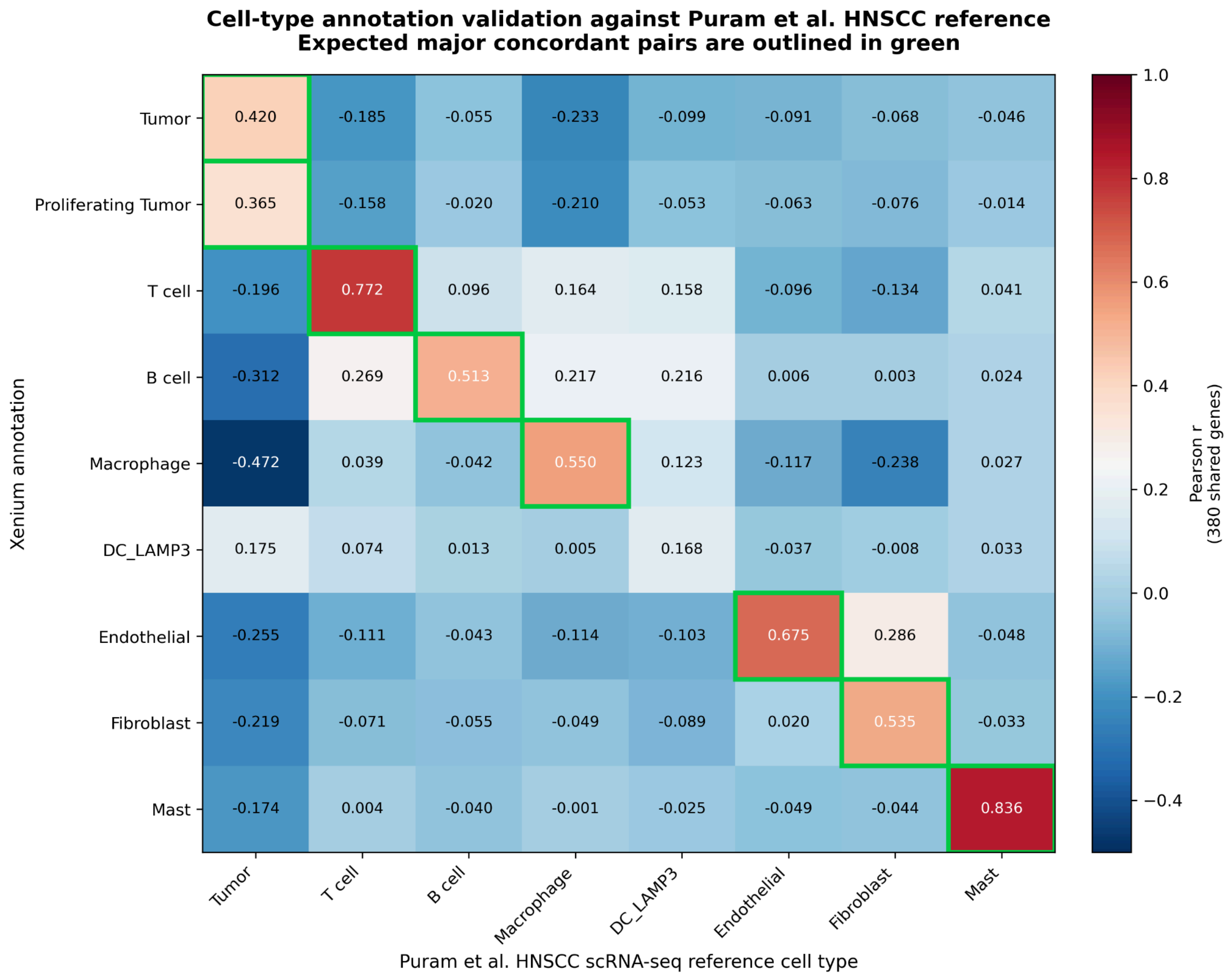


**Supplementary Figure S3. Cell-type annotation validation against the Puram et al. HNSCC single-cell reference.**

Pearson correlation was calculated between mean expression profiles of Xenium annotated cell types and Puram et al. HNSCC scRNA-seq reference cell types using 380 shared genes. Expected concordant cell-type pairs are outlined in green. Most major tumor, immune, and stromal annotations showed expected concordance with the reference dataset. DC_LAMP3 showed weaker correlation with the reference dendritic cell category, consistent with differences in dendritic cell state resolution across datasets, and was further validated using curated mature/migratory dendritic cell marker scoring.

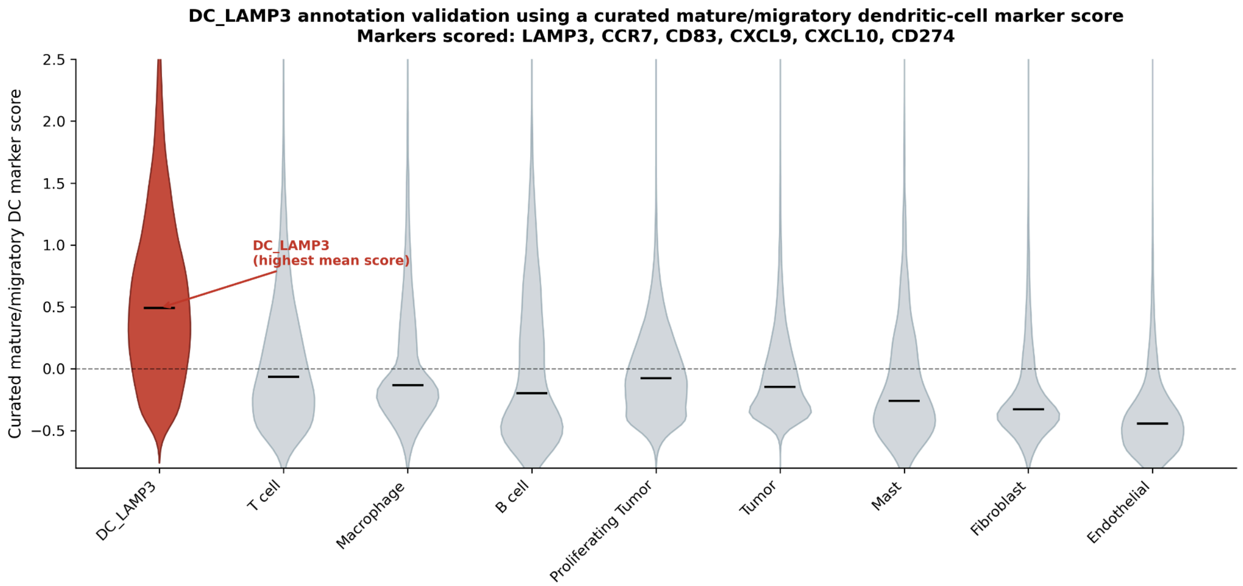


**Supplementary Figure S4. DC_LAMP3 annotation validation using a curated mature/migratory dendritic-cell marker score.**

Cells were scored using a curated mature/migratory dendritic-cell marker set consisting of LAMP3, CCR7, CD83, CXCL9, CXCL10, and CD274. The DC_LAMP3 cluster showed the highest marker score across annotated cell types, supporting its annotation as a mature dendritic-cell population.
